# Genetic Continuity of Indo-Iranian Speakers Since the Iron Age in Southern Central Asia

**DOI:** 10.1101/2021.11.04.466891

**Authors:** Perle Guarino-Vignon, Nina Marchi, Julio Bendezu-Sarmiento, Evelyne Heyer, Céline Bon

## Abstract

Since prehistoric times, South Central Asia has been at the crossroads of the movement of people, culture, and goods. Today, the Central Asia’s populations are divided into two cultural and linguistic groups: the Indo-Iranian and the Turko-Mongolian groups. Previous genetic studies unveiled that migrations from East Asia contributed to the spread of Turko-Mongolian populations in Central Asia and the partial replacement of the Indo-Iranian population. However, little is known about the origin of the latter. To shed light on this, we compare the genetic data on two current-day populations– Yaghnobis and Tajiks – with genome-wide data from published ancient individuals. The present Indo-Iranian populations from Central Asia display a strong genetic continuity with Iron Age samples from Turkmenistan and Tajikistan. We model Yaghnobis as a mixture of 93% Iron Age individual from Turkmenistan and 7% from Baikal. For the Tajiks, we observe a higher Baikal ancestry and an additional admixture event with a South Asian population. Our results, therefore, suggest that in addition to a complex history, Central Asia shows a remarkable genetic continuity since the Iron Age, with only limited gene flow.

## INTRODUCTION

Central Asia is a large region stretching from the Caspian Sea in the west to Lake Baikal in the east, encompassing Tajikistan, Kazakhstan, Turkmenistan, Uzbekistan, Kyrgyzstan and north Afghanistan. This region has found itself at the crossroads of migration routes since modern humans left Africa^1,2^, leading to a long-term presence of humans, a rich history, and a high cultural diversity. For illustration, agropastoral communities present since Djeitun culture^3^ 6000 years BC were replaced during the Chalcolithic (4800-3000 BC) by the emergence of denser villages and the premises of irrigated agriculture. During the Middle Bronze Age, the Bactrio Margian Archaeological Complex (BMAC) civilization flourished in southern Central Asia^4^ with characteristic proto-urban cities, powerful irrigation techniques, and a marked social hierarchy^4^. A pastoral nomadic lifestyle emerged later in northern Central Asia around 3000 BC and gained importance in this region during the late Bronze Age (2400-2000 BC). At the end of the Bronze Age, from about 1800 BCE, the Oxus civilization during its final phase underwent important transformations: while remaining in the same tradition, the material culture was impoverished with some ceramic forms and artifacts disappearing; some habitat sites were abandoned, monumental architecture disappeared, the level of technological development seemed to decrease^5^; international trade, which had been flourishing during the previous peak phase, slowed down considerably, or even came to a halt, except for contacts with the steppes of northern Central Asia^6^; funerary practices changed with the appearance of new modes of burial, before the total disappearance of burials during the Early Iron Age, that can be linked to an ideological evolution^7^. The period between 1800 and 1500 BCE saw Andronovo-like culture take over, until the rise of Yaz culture^8,9^. Then, Central Asia was the scene of the eastwards conquests of Achaemenids, Greeks, Partho-Sassanians and Arabic people and of the westward movement of various Asian peoples like the Huns, the Xiongnus, and the Mongols^10^, before being a trade centre along the Silk Road, particularly during the Sassanid Empire and after the Islamic invasion.

Today, the complex demographic history of Central Asia results in a composite genetic diversity, with modern Central Asian populations being divided into two culturally distinct groups: a first group composed of Turkic and Mongolic-speaking populations (referred to later as Turko-Mongol populations including Kyrgyz, Kazakhs...), who are semi-nomadic herders^10^ and show genetic affinities with Eastern Asian and Siberian populations; and a second group formed by Tajiks and Yaghnobis who live in southern Central Asia, speak Indo-Iranian languages, practice agriculture, are sedentary and who are genetically more similar to present-day western Eurasian populations^2,11^ and Iranians^12^. Moreover, Yaghnobis are known to have been isolated for a long time with no evidence of recent admixture^12^. Modern DNA studies suggested that the Indo-Iranian group was present in Central Asia before the Turko-Mongol group^11^, maybe as early as Neolithic times; the Turko-Mongol group emerged later from the admixture between a group related to local Indo-Iranian and a South-Siberian or Mongolian group^11,13,14^ with a high East-Asian ancestry (around 60%). Turkmens, however, genetically stand out from the Turko-Mongol group, being intermediate with the Indo-Iranian group^15^, which suggests a recent language and culture shift^16^, possibly through a mostly elite dominance-driven linguistic replacement.

Paleogenetic studies confirmed that multiple migration waves and admixture events, in which steppe populations played an important role, have occurred in Eurasia in the last 10,000 years^13,17–20^. Although the settlement of Europe was extensively studied^21–26^, there have been few studies exploring the population history of Central Asia, and even fewer focusing on southern Central Asia. In northern Central Asia (Kazakhstan, Southern Russia), genetic studies evidenced eastward and westward movement of populations since the late Neolithic period^13,17,18,27,28^, leading to a gradient of western steppe genetic ancestry. In southern Central Asia where most of the ancient genomes date back to the late Neolithic and the Bronze Age, it was shown that populations from the BMAC were strongly related to southern Iranian ancient populations with some individuals displaying additional steppe-ancestry^18^.

However, the relation between modern Indo-Iranian speaking populations and ancient populations from southern Central Asia remains unclear: what are the genetic sources of modern Indo-Iranian speakers? Can they be traced back to the Iron or the Bronze Age? Is there one or several different population histories among a given linguistic group of populations? What is the role of the Turkmens in this story?

Paleogenetic studies brought additional tools to seek the origins of these populations. To explore the origins of modern Indo-Iranians in relationship with their Turko-Mongol neighbours, we jointly analyzed genome-wide data in 16 modern populations (one Yaghnobi and four Tajik populations, 11 from distinct Turko-Mongol ethnic groups from Central Asia, i.e. in Uzbekistan, Kyrgyzstan, Tajikistan and from West Mongolia and South Siberia) as well as 1501 present-day genomes from Eurasia and Africa^29,30^ and 3109 ancient published genomes from all Eurasia^13,17,31–40,18,41–45,19,20,22–24,27,28^ (Table S1), including 126 ancient genomes from southern Central Asia^17,18^ (fig1.a).

**Figure 1.**
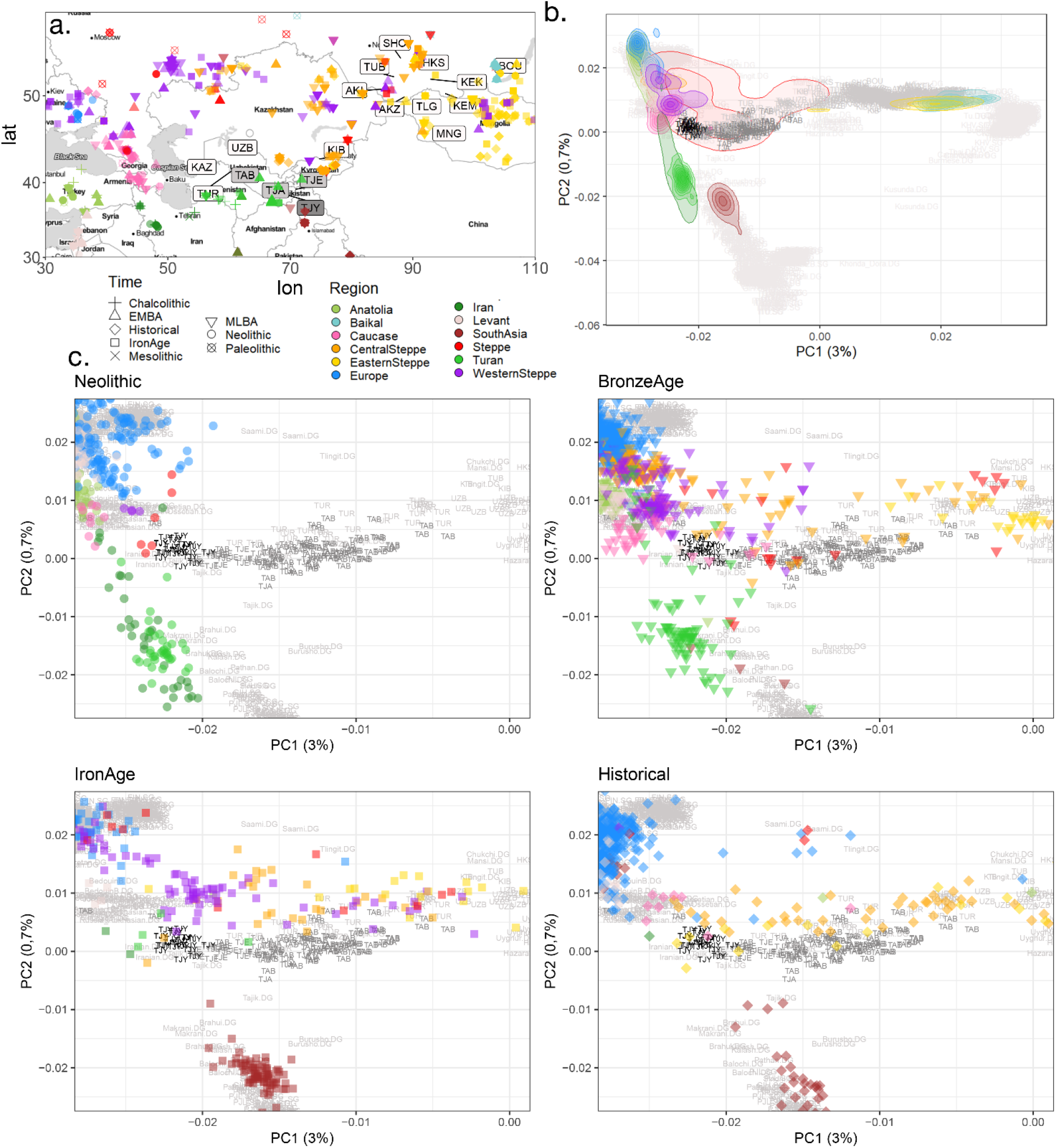
Geographic and genetic structure of our dataset a – Map of the published ancient samples in our dataset b – PCA computed on a set of 236,566 SNPs for present-day Eurasians populations including 527 present-day Central Asia individuals genotyped on an 300k SNPs array^15^ and we projected the 3102 ancient genomes onto the PCs. Ancient genomes are represented with different colors by region, with density line to facilitate the reading c – Details of b. focusing on Indo-Iranians individuals with ancient individuals from Neolithic, Bronze Age, Iron Age and Historical times.

## RESULTS

### Modern Indo-Iranian genetic affinities with ancient samples

To explore the relation between present-day Central Asian individuals and the Eurasian genomic diversity, ancient and modern, we first performed a Principal Component Analysis (PCA) (fig1.b) on 1915 modern genomes and projected 3102 ancient genome-wide data onto it. Regarding the present-day Eurasian diversity, the three top PCs roughly mimic the geographical repartition of modern populations: the PC1 (3% of variance) discriminates between Eastern and Western Eurasian individuals, the PC2 between South Asian and modern European individuals, and the PC3 discriminates against the Baikal populations from the East-Asian cluster (see Supplementary fig. S1). Present-day Indo-Iranian individuals from Central Asia cluster together on the first three PCs while Turko-Mongol individuals form a gradient from the Indo-Iranian cluster to ancient Baikal samples on PC3, in agreement with cultural categorization instead of geography. However, a substructure appears within the Indo-Iranian group with the Yaghnobis (TJY) falling closely to the Western cluster, while the Tajiks populations (TJA, TJE, TAB) stretch toward the Baikal cluster, indicating some additional East Asian or Baikal Hunter-Gatherer (BHG) proximity.

For the ancient individuals, Bronze Age, Iron Age, and historical Steppe individuals fall on a cline stretching up from European to East Asian groups, with *Western_Steppe* individuals clustering on the bottom of the European cluster and *Central_Steppe* individuals spreading from the *Western_Steppe* cluster to the *Okunevo_BA* cluster close to Baikal and Siberian modern individuals. The ancient individuals of southern Central Asia (Neolithic, Bronze Age and Iron Age) follow a cline stretching from Neolithic Iranian individuals *(Iran_N)* to present-day Iranians and Yaghnobis.

Contrastingly, the Iron Age samples (*Turkmenistan_IA* and *Ksirov_Kushan* individuals) are located close to modern Indo-Iranian populations, although slightly negative values on the first axis and positive values on the third axis suggest an addition of Baikal ancestry in the present-day Indo-Iranians. Finally, it appears from this PCA (fig 1.c) that ancient and present-day Indo-Iranian populations from Central Asia form together a cline between Iranian Neolithic farmers and *Central_Steppe* Bronze Age, with a clear shift in ancestry toward Steppe between Bronze Age and Iron Age as observed before^18^, and a smaller shift toward eastern Asian ancestry between Iron Age and present-day. This shift is more pronounced in Tajiks than Yaghnobis.

To confirm our initial observations and identify genetic structures, we performed an unsupervised clustering using ADMIXTURE^46^ on the same dataset used for PCA (see Supplementary fig. S2). Consistently with the PCA, we evidenced in all modern Indo-Iranians the presence of a genetic component maximized in Iran Neolithic farmers (dark green; mean value for Yaghnobis: 37%; 25% for Tajiks), of another maximized in EEHG (Eastern European Hunter-Gatherer) and WSHG (Western Scandinavian Hunter-Gatherer) (pale green, mean value for TJY: 13%, mean value for Tajiks: 10%) and of a third component (dark blue; mean value for TJY: 36%; for Tajiks: 29%) that is not completely maximized in any population of our dataset, but is found in present-day Europeans and in Anatolian Neolithic farmers (*Anatolia_N*). In addition, a fourth component maximized in BHG *(Shamanka_EN)* and largely present in all modern Turko-Mongol populations (red; 50% on average) is also inferred to a lower extent in the modern Indo-Iranian populations, with a significantly smaller proportion in Yaghnobis than in Tajiks (mean value respectively 7% and 14%; t-test p-value = 2.10^-16^). Finally, the Tajiks present a small proportion (4%) of modern East Asian ancestry (pink component, maximized in Han population), which is largely present in all Turko-Mongol populations from Central Asia (mean value 10%), and around 8% of the component maximized in present-day South-Asian populations (orange), which are both absent in Yaghnobis.

ADMIXTURE analysis is also congruent with PCA concerning the ancient groups (see Supplementary fig. S3 and S4). Indeed, Iron Age southern Central Asia individuals present a remarkably similar profile to Yaghnobis’ profile: for instance, the individual labelled as *Turkmenistan_IA* has a profile with about 25% of WSHG/EEHG component, 30% of *Iran_N* component and 35% of the Anatolian farmer ancestry component but missing BHG ancestry (fig. 2). Bronze Age Central Steppe pastoralists show a similar profile except for a significant increase in Iranian ancestry, and Western Steppe pastoralists have the beige component maximized in WEHG (Western European Hunter-Gatherer), which is absent in modern Indo-Iranian populations.

**Figure 2.**
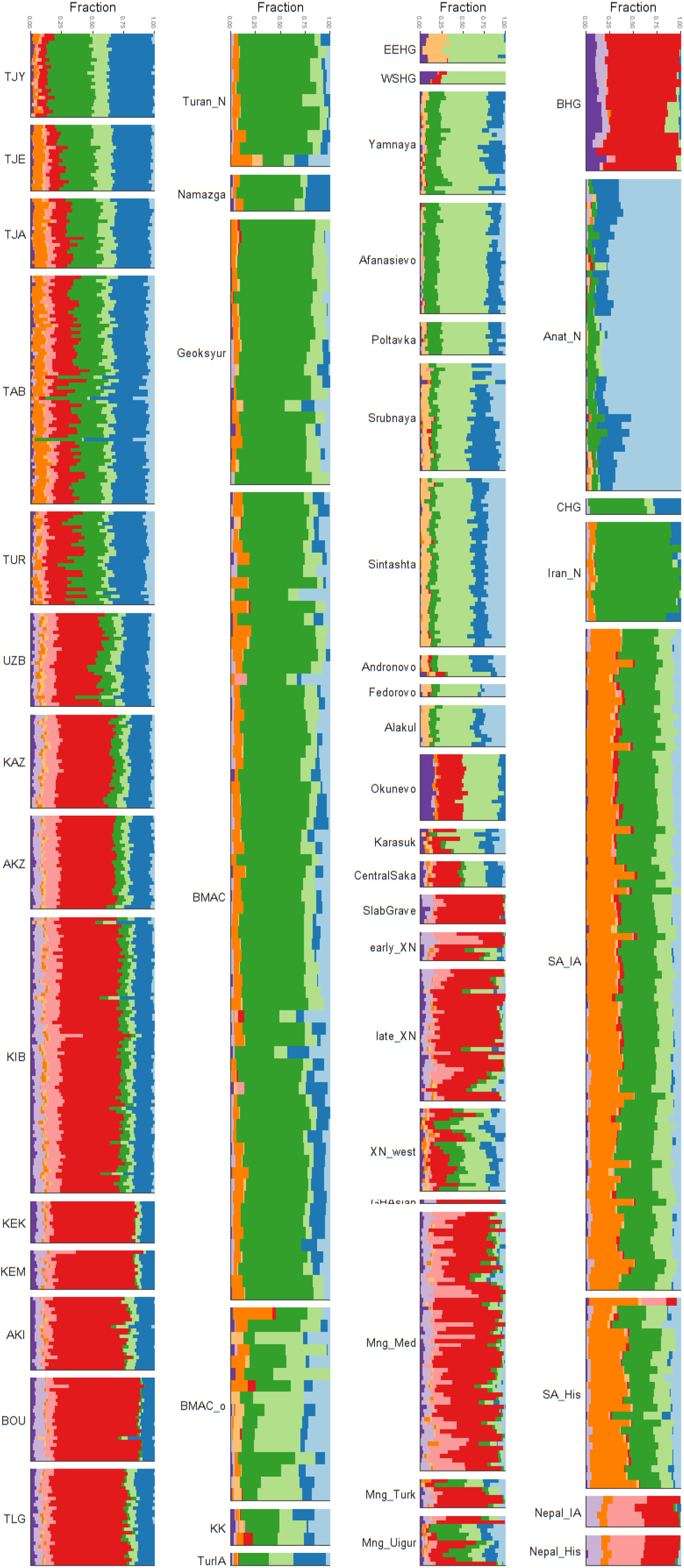
ADMIXTURE analysis of 5019 individuals (3102 ancient and 1915 modern). The results for a subset of the dataset (present-day Indo-Iranian individuals and ancient populations discussed in the main text) are displayed for K=10 clusters, which has the lowest cross validation value (0.994). The full analysis is shown in SI. In the first column, are the modern individuals from Central Asia, second column the ancient individuals from southern Central Asia, third column ancient individuals from the Steppe, and in the last column are miscellaneous individuals discussed in the main text.

Thus, modern Indo-Iranian speaking populations appear as midway between Central Steppe and southern Central Asia Bronze Age populations, quite similarly to the Turkmenistan Iron Age individuals, with a limited **impulse** from eastern Asian and southern Asian groups.

### Population continuity within the Indo-Iranians

To formally test for genetic continuity with Iron Age southern Central Asia and the limited admixture with Baikal-related populations at the source of the present-day Indo-Iranian speaking populations, we performed D-statistics, f3-statistics and qpAdm modelling on the same dataset used for PCA et ADMIXTURE analysis as well as on a dataset formed by shotgun sequences from 3 Yaghnobis (TJY), 19 Tajiks (TJE) and 24 Turkmens (TUR)(reference to be given) as well as the ancient genomes for a final set of ~700k SNPs.

We identified and characterized gene flows that occurred since the Iron Age by computing D-statistics of the form D(Mbuti, *Ancient population*; *Turkmenistan_IA,* present day Indo-Iranian) for every ancient population in our dataset (fig. 3, Table S5). These statistics are expected positive when gene flows occurred from the *Ancient population* to the present-day Indo-Iranians. For the Yaghnobis, only one individual, an Iron Age individual from Nepal genetically close to East Asian populations (Nepal_Chokhopani_2700BP.SG)^45^, has a significantly positive D-statistic (Z>3). Tajik individuals (TJE) display a higher number of ancient populations (N=41) whose D-statistic is positive; the common characteristic of these ancient populations is to exhibit a large amount of BHG ancestry, consistently with the ADMIXTURE analysis (fig 2). We also note that the Tajiks present a positive D-statistic with a historical individual from India (Great Andaman) (fig 3) showing a possible connection with South Asia. Thus, modern Indo-Iranian populations descend from groups related to those present in Turkmenistan as early as Iron Age, with a contribution from another East Asian population who brought the BHG ancestry and, except for Yaghnobis, a contribution from a South Asian population.

**Figure 3:**
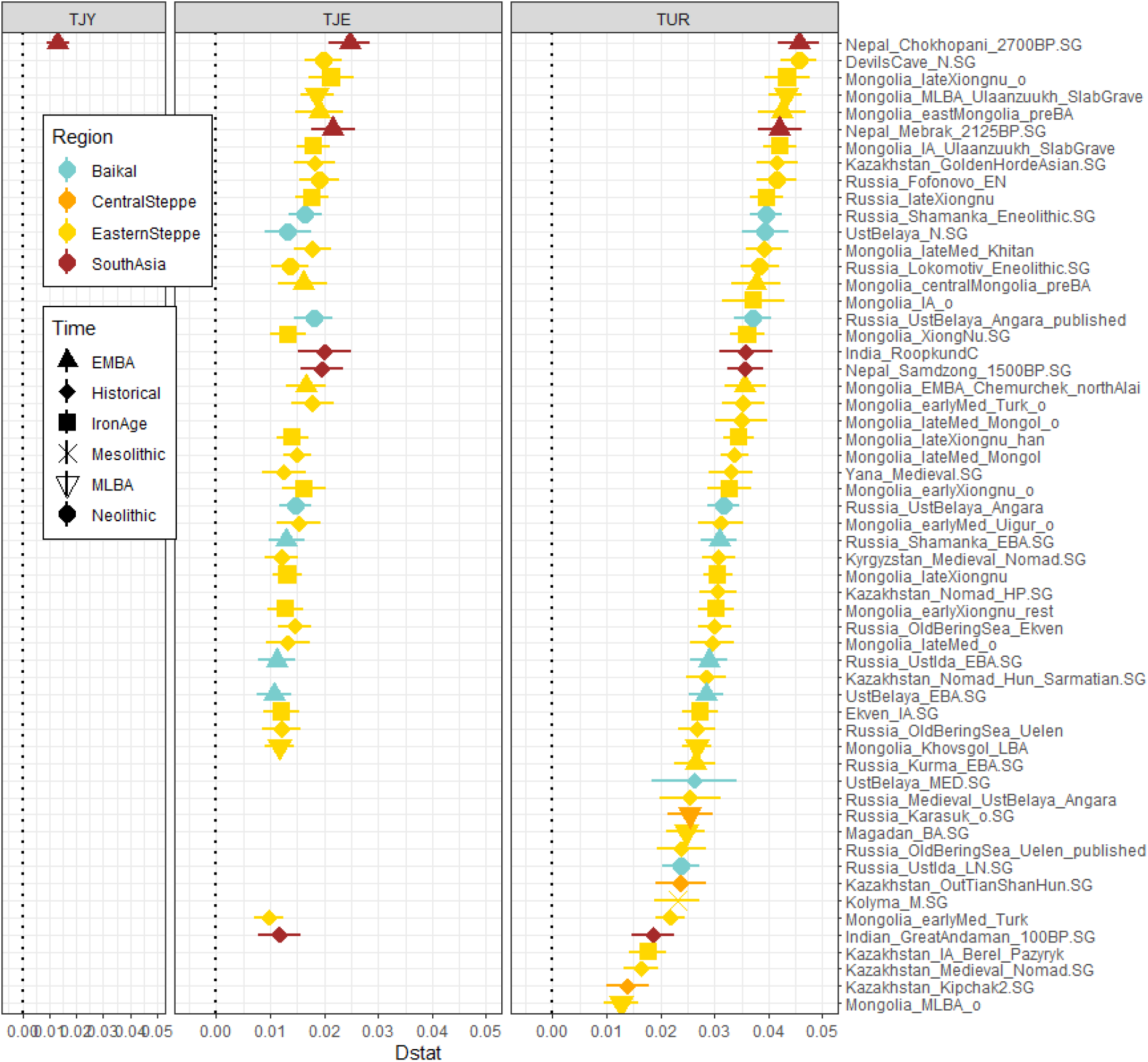
Gene flow in Indo-Iranian populations since Iron Age. Positive D-statistics (Z>3) of the form D(Mbuti, *Ancient population*; *Turkmenistan_IA*, TJY/TJE/TUR). A positive D-statistic demonstrates that a gene flow occurred from the Ancient population to the Indo-Iranian population or Turkmen population compared to *Turkmenistan_IA.* The estimated statistic ± 3 standard errors is indicated.

Then, we formally test if the contributions detected with D-statistics are due to admixture events that occurred since the Iron Age. We first computed f3-statistic^47^ of the form f3(TJY/TJE *; Source1*, *Source2*), that is expected to be negative (Z<-3) if the Indo-Iranian populations can be modeled as an admixture between the two sources (Table S7). Only combinations implying a population from East Asia ancestry (like XiongNu) and westerner populations representing the components seen in the ADMIXTURE analysis (Iranian Neolithic, Anatolian farmer, and Steppe ancestry) were significant (fig. 2). These statistics attest to the existence of an actual admixture between a population probably presenting a mix of Iran Neolithic, BMAC, Anatolian early neolithic and Bronze Age Steppe ancestry with a population with a strong affinity to the *Baikal_HG* ancestry. Yaghnobi population has significantly fewer pairs with a negative f3-statistic than the 3 Tajik populations, probably due to their long-term isolation. We also specifically calculated f3-statistic of the form *f3*(*Ancient population, Turkmenistan_IA*; TJY/THE/TJA/TUR) and obtained several negative f3-statistics always with the same ancient populations implied in the positive D-stat (see Supplementary fig. S2, Table S6) showing that Indo-Iranians can be successfully modelled as the admixture of Iron Age Turan and BHG-related population.

We then modelled Yaghnobi and Tajik populations using *qpAdm*^23^ to estimate mixture proportions. To test which proximal populations fit the best in our model, we used the rotating method^23^ and we excluded all combinations with a *p-value* ≤ 0.01. We first tried a two-ways admixture testing several possibilities among rotating sources. For the Yaghnobis, the only model retained was the one with ~93-88% from *Turkmenistan_IA* and ~7-12% ancestry from XiongNu. With 3-ways modelling, we could not reject different models for TJY: 3 models imply 90% ancestry from *Turkmenistan_IA* and 7% ancestry from XiongNu, and around 3% of ancestry from *Europe_EN,* BMAC or *Ukraine_Scythian*; we also obtained a model with *Ukraine_Scythian,* BMAC and XiongNu inferring the older admixture at the origin of *Turkmenistan_IA*. When testing for more admixture sources, we obtained only two 4-ways models and one 5-ways model (Supp. Data). One interesting model is a 4-ways model with 17% Ukrainian Scythians, 60% *Turkmenistan_IA*, 14% BMAC and 8% XiongNu, i.e. this model shows a close affinity of Yaghnobis with Western Steppe-like populations.

To model Tajiks, all 2-ways admixture models were excluded and we obtained one 3-ways admixture model *(p-value* = 0.49) implying around 17% ancestry from XiongNu, almost 75% ancestry from *Turkmenistan_IA,* and around 8% ancestry from a South Asian individual (*Indian_GreatAndaman_100BP*)^48^ representing a deep ancestry in South Asia.

Thus, the *qpAdm* modelling shows that at least 90% of the ancestry of current Indo-Iranian ancestry is modelized as inherited from Iron Age individuals from southern Central Asia with an affinity with BMAC. Consequently, Indo-Iranians present a strong genetic continuity in the region since the Iron Age with anecdotic admixture with BHG ancestry related individuals, and, for the Tajiks, with South-Asian ancestry related populations possibly after Iron Age.

Finally, we used *DATES*^18^ to estimate the number of generations since the admixture events. We obtained 35±15 generations for the admixture between *Turkmenistan_IA* and XiongNu-like populations at the origins of the Yaghnobis, i.e. an admixture event dating back to ~ 1019±447 years ago considering 29 years per generation^49^. For Tajiks (TJE, TJY, TJA) we obtained dates from ~ 546 ±138 years ago (18.8± 4.7 generations) to ~ 907 ± 617 years ago (31.2 ± 21.3 generations) for the West/East admixture. We also obtained a date of ~ 944 ±300 years ago for the admixture with the South Asian population.

### Iron Age Turkmenistan ancestry

Previous studies^13,18^ have already shown *Turkmenistan_IA* can be modelled as an admixture between BMAC and some steppe populations, and on the PCA (fig 1.c), *Turkmenistan_IA* indeed belongs to the Steppe cline. However, the steppes are split between several groups (Western steppe, Central Steppe, Eastern Steppe) depending on their amount of Eastern Asian ancestry. The ADMIXTURE analysis discriminates the Western and Central steppe ancestries by the presence of a red and mauve component (maximized respectively in East-Siberia and East-Asia populations) in the latter, which is absent from *Turkmenistan_IA*, indicating an affinity with the Western Steppe. Nevertheless, we noted that Andronovo or Sintashta individuals also lacked this component while being classified as Central_Steppe. Thus *Central_Steppe* group is highly heterogenous and gathers populations with some East-Asian ancestry like Karasuk or Central Saka and others more Western Steppe-like as Andronovo and Sintashta. Furthermore, we obtained the higher f3-outgroup statistic of the form f3(Mbuti; Ancient pop, *Turkmenistan_IA*) for ancient populations from BMAC complex or West Eurasia, highlighting the double origin and affinity with the West. This affinity is further confirmed with D-statistics of form D(Mbuti, *
Turkmenistan_IA*; *Western_Steppe, Central_Steppe*) (fig 4.B) that are significantly negative (Z<-3) when a *Western_Steppe* population is opposed to a *Central_Steppe* population with an East-Asian ancestry, like Central Saka or Karasuk (fig 4.B). With D-statistic of the form D(Mbuti, *Turkmenistan_IA*; HG1, HG2) – HG1 and HG2 belonging to WEHG, EEHG, WSHG, and BHG populations – we evidenced that the Steppe populations admixed with BMAC lacked East Asian or Baikal component (fig 4.A). Indeed, we only see significant D statistics when BHG was confronted with the other HGs (fig 4.A). Using HG populations avoids inferences from recent admixture; nevertheless, it failed to discriminate between most of the different Steppe groups of this period at this level. This suggests that *Turkmenistan_IA* is devoid of the East Asian ancestry observed in several Central steppe groups as early as Bronze Age.

**Figure 4:**
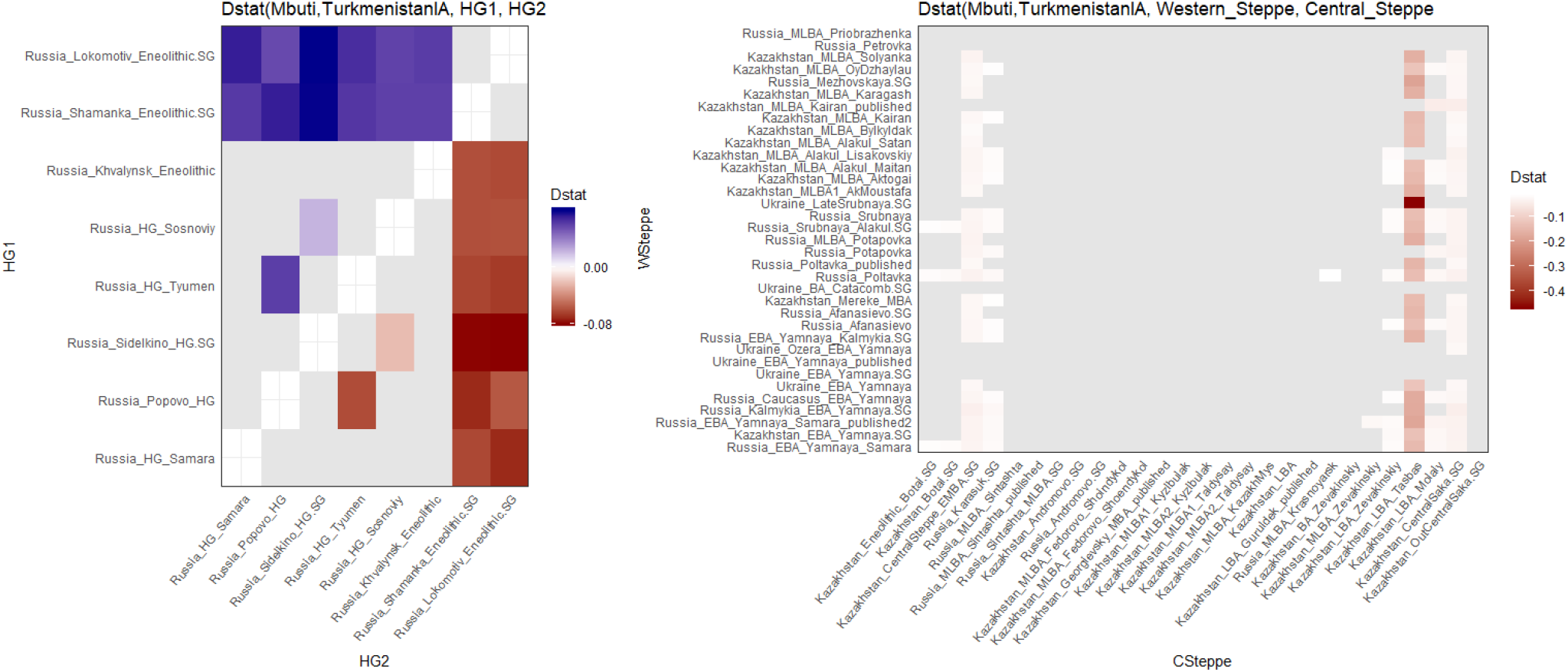
Absence of affinity of *Turkmenistan_IA* with East Asia ancestry shown by D-stat. In grey are non-significative (Z<3) D-statistics, in blue significative positive D-statistics and in red significative negative one. Only populations with strong East-Asian or BHG ancestry show a significative D-statistic.

Finally, we tested different Steppe populations which admixed with BMAC to model *Turkmenistan_IA* with *qpAdm.* We first constituted a set with Poltavka, Srubnaya *Western_Steppe*) and 4 individuals from Russia labelled as Andronovo (*Central_Steppe*)^50^, to estimate the affinity with Europe and Western Steppe previously highlighted with D-statistics and f3-statistics. We only obtained one model with 2 sources that we could not exclude, and it implies an admixture of 43% BMAC and 57% Andronovo *(p-value* = 0.31) suggesting that Andronovo individuals are the best proxy for the steppe population which admixed with BMAC to form the Iron Age southern Central Asia group. When testing for the best model between Andronovo and Karasuk (Central Steppe with East Asian component) to estimate the affinity with Asia, we produced a single fitting/ relevant model implying Andronovo (p-value = 0.51) with roughly the same proportions. Further tests explored the best model between Andronovo and Sintashta, two genetically close populations, and the single significantly outcome was the one with Andronovo and BMAC (*p-value* = 0.498) in the same proportions. Eventually, we tested the best model between the individuals labelled Andronovo and two populations belonging to the Andronovo-complex: Fedorovo Shoindykol^18^ and Alakul Lisakovskiy^18^. Once again, the only valid model was the one with Andronovo and BMAC. Overall, we can say that the Iron Age population from southern Central Asia emerges from the admixture of BMAC with a Bronze Age population close to the Andronovo individuals, which presents a profile with an affinity with Western Steppe rather than with a Central Steppe with an affinity with East Asia (like Karasuk).

### A Turkmens’ history

Despite speaking a Turko-Mongol language and having the same cultural practices as other Turko-Mongol ethnic groups^51^, Turkmens are genetically closer to Indo-Iranian populations than to Turko-Mongols^52,53^.

Indeed, Turkmens (TUR) fall into the Tajiks cluster and not in the Turko-Mongol cline in the PCA (fig. 1) and in the ADMIXTURE analysis (fig. 2), all Turko-Mongol populations from Central Asia except Turkmens show a significant (t-test, *p-value* < 2.10^-16^) high amount of Baikal (red component, mean 50%) and East-Asian ancestry (pink component, maximized in Han population). Turkmens, for their part, display a completely different pattern, with an amount of Baikal component (mean value: 22%) closer to the proportion in Tajiks (mean value: 15%) and almost no East-Asian component. They do not show as much South-Asia related ancestry as Tajiks, suggesting that the admixture with South-Asian populations occurred or continued after Turkmens split from the remainder of the Indo-Iranian group.

We have established genetic affinity profiles with ancient populations for all Central Asia populations including Turkmens of the first dataset, based on f3-outgroup statistics of the form f3(Mbuti; Ancient pop, Present-day pop) (fig. 5; Table S8). The f3-outgroup values comparing Turkmen to any ancient population are strongly correlated with the one comparing Tajiks to any ancient populations (fig 5.a). On the other hand, f3-outgroups values comparing Eastern steppes and Baikal groups to a Turko-Mongol population (Kazakhs) are higher than those comparing these ancient populations to Turkmens (fig 5.b). The Turkmens are more similar to Indo-Iranian populations than to any Turko-Mongol population on the amount of shared Siberian/East Asian ancestry.

**Figure 5:**
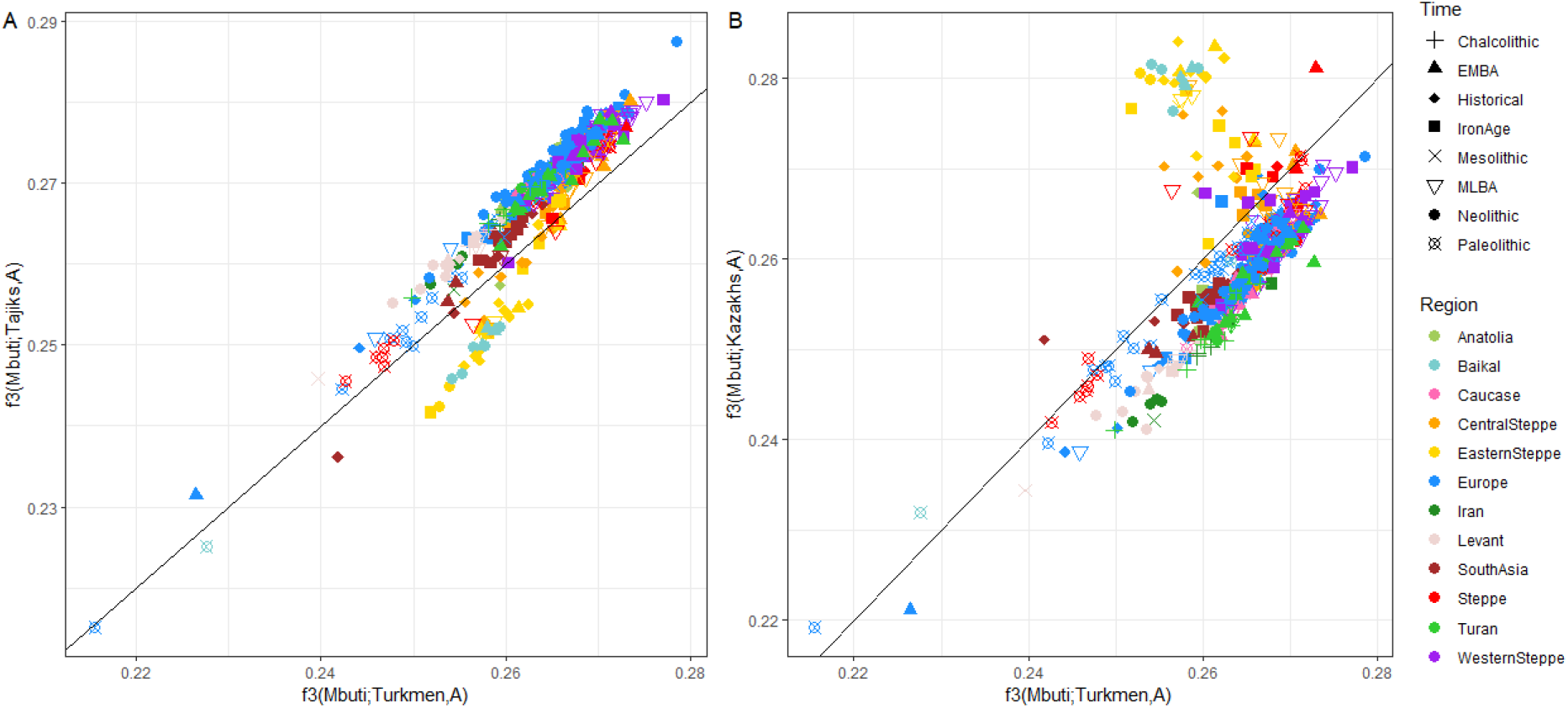
Turkmens’ affinity with Tajiks rather than with Turko-Mongol groups shown by f3 statistics of the form f3(Mbuti; TUR/TJA/AKZ, *Ancient population*) (A) Outgroup f3-statistics for Turkmen and for Tajiks (TJA) plotted against each other (B) Outgroup f3-statistics for Turkmen and for Kazakhs (AKZ), belonging to the Turko-Mongol group, plotted against each other

Finally, we modeled Turkmens as a mixture of Central Asia basal ancestry (represented by Yaghnobis) and East-Asian ancestry (we obtained a negative value for f3(TUR; TJY, *DevilsCave_N);* f3 = −0.0025, Z =-5.266). *qpAdm* modelling for Turkmens produces a single nonrejected model *(p-value* = 0.048007) implying 6% of Golden Horde Asian and 94% of Tajiks (TAB) (with TJY, XiongNu, GoldenHordeAsian, TAB, Turkmenistan_IA as potential rotating left population). For this admixture event, we estimated a date of 687 ± 100 BP (23,7 ± generations) with *DATES.*

These results enlighten that Turkmens were an Indo-Iranian-like population not so long ago, who recently shifted language and culture without a substantial genetic change in population.

## DISCUSSION

Our research provides insight into the history of Indo-Iranians by using evidence to trace modern populations back to the Iron Age in southern Central Asia. As proposed by former genetic studies^2,11^ and as supported by historical^54^ and archaeological evidence^55^, we found that Indo-Iranian speakers settled in Central Asia long before Turko-Mongol speakers^11^. The main event at the bottom of Indo-Iranian ancestry in southern Central Asia occurred at the end of the Bronze Age/Early Iron Age, through the admixture between local BMAC groups and Andronovo-related populations perhaps linked to the end of the Oxus Civilization. We note here that the steppe group who admixed with BMAC did not present East Asian ancestry, which is consistent with both the archeological^56^ and genetic^38^ findings of the East Asian ancestry arriving in the Central Steppe core only at the end of the Iron Age.

The populations falling under the name Andronovo form a complex group. Indeed, when screening the individuals used under the label Andronovo in our dataset, we note that they all belong to one site, Kytmanovo^50,57^, which is eastward, but show a genetic profile very close to the Sintashta individuals, whose area expanded near the Caspian Sea. Individuals from other cultures belonging to the Andronovo complex have been sequenced^17,18^ but overall they form a moderately heterogenous genetic group. Moreover, some studies have shown that Steppe groups can be labelled similarly but be different genetically, such as, for instance, Srubnaya Alakulskaya individuals being closer to Andronovo individuals than to Srubnaya from the Samara region^28^. The nomadic populations from the end of the Bronze and Iron Age being very genetically heterogenous, we suspect that the source of the western steppe ancestry found in Iron Age southern Central Asia may not be sampled yet. It is interesting to notice that the gene flow between the Steppe and southern Central went two-ways^38,58^. A recent study from Gnecchi-Ruscone *et al.*^58^ has highlighted that a gene flow from BMAC contributed to the genetic formation of Scythians. Our findings combined with these studies strongly corroborate the hypothesis based on archaeological evidence that southern Central Asia civilizations since BMAC and western Steppe culture had a strong cultural connection^6,8,59–62^.

Overall, we demonstrate here a remarkable example of genetic continuity since the Iron Age in Indo-Iranian populations from Central Asia despite the frenzy of population migrations in the area since the Bronze Age. Similar to Zhabagin *et al.* work^63^, the present study shows no impact of the Arab cultural expansion in Central Asia on the Indo-Iranian speaker’s genetic diversity, despite the first one leading to a shift in language for Tajiks. We also do not see a gene flow from Iran despite the Persian cultural expansion which led to a language shift from an east-Iranian language to a west-Iranian in Tajiks – when Yaghnobis kept their East-Iranian language^64^.

Yaghnobis, for their pair, are characterized by strong genetic stability over time (small amount of negative admixture f3-statistics, fewer significative D-statistics), which can be linked back to their long-term isolation^12,65^. Yaghnobis are indeed an isolated ethno-linguistic population historically present in the hardly accessible valley of the Yaghnob River. Evidence suggests that the separation between Yaghnobis and Tajiks occurred at least 1000 years ago, which explains the high genetic differentiation observed in Indo-Iranians by previous studies^51,65^. Interestingly, it implies that Yaghnobis could represent a good proxy for the ancestry present in Central Asia before the migration waves that led to the current genetic diversity, despite the strong drift that occurred.

The amount of East-Asian ancestry due to admixture with modern Turko-Mongol groups remains low even in Tajiks, consistent with the findings of Martinez-Cruz *et al.*^2^, who observed the light impact the westward invasions (Huns, Mongols) had on Indo-Iranian groups in Central Asia. On the other hand, we have highlighted for Yaghnobis, Tajiks, and Turkmens a small amount of gene flow from BHG-ancestry dating to around 1000 years ago, suggesting a recent wave of westward migration from the Altai mountains, after the Iron Age. This recent wave can be linked to the origin of the Turko-Mongol in Central Asia which has been demonstrated by Martinez-Cruz *et al.*^2^ and Li *et al.*^66^ to be from an ancestral group of Turkic speakers from the Altai region. Our quite recent date of admixture differs significantly from the date obtained by Palstra *et al.*^11^ which placed the admixture event back to 8 ky BP for Tajiks and 2.3 ky BP for Kyrgyz. The more recent inferred dates of admixture for Tajiks compared to Yaghnobis could be explained by the fact that Tajiks received a more continuous gene flow from the eastward source. Continuous gene flow that occurred after the first admixture event that formed the Yaghnobis genetic composition. Indeed, the *qpAdm* method cannot detect a continuous admixture which can be expected in this context. Furthermore, the search of their ancestry confirms a genetic homogeneity within Yaghnobis, Tajiks, and Turkmens, despite their cultural, notably linguistic differences, with some genetic differences emerging from various patterns of gene flow in Tajiks and Turkmens.

Notably, we evidenced an admixture event from South-Asia restricted to the Tajik population, undocumented before despite evidence in Iranian Turkmens^67^. According to previous archaeological studies^68,69^, multidirectional cultural exchanges with South Asia are known to have taken place as early as the Chalcolithic period: notably from Sialk culture and other Iranian cultures towards Balochistan^68^ or from Geoksjur culture of Turkmenistan to southern Afghanistan. In the opposite direction, from south to north, Mundigak III type ceramics find parallels as far as Badakhshan in northeast Afghanistan, material from Balochistan and shells used in necklaces and bracelets from the Arabian Sea are found at the Sarazm site in Tajikistan, showing a long-distance commercial exchange. All these ancient populations were on the move with probably quite frequent exchanges and cultural blends between populations, Iron Age included^69^. Intriguingly, genetic proximity between southern Central Asian and South Asian groups has already been suggested for BMAC samples^18^ and raises the question of the timing of this gene flow. Two models can be considered: the first one assumes the formation of a homogeneous basal Indo-Iranian background (as observed today in Yaghnobis) and subsequent recent gene flow from South-Asian populations; the second model acknowledges the presence of South-Asia ancestry in some Bronze Age BMAC samples^18^ and suggests Tajiks and Yaghnobis could have derived from distinct BMAC populations, respectively with and without South-Asian ancestry, who have both experienced independent admixture with Andronovo-like steppe populations during Iron Age, and eastern nomads with BHG ancestry afterwards. Because the date of the gene-flow from South Asian populations in Tajik genomes is relatively recent, the data favours the first hypothesis; however, uncertainties on the model of admixture (one *versus* several pulses) may be compatible with continuous gene-flow since the Bronze Age. Additionally, our recent date of admixture fits with the arrival of the South-Asian ancestry at the same that the shift from east to west-Iranian language in Tajiks linked to the Persian expansion 1500 years ago^64^.

Lastly, the case of Turkmens is a notable example of a population changing language and cultural practices without substantial changes in their genetic ancestry. Indeed, Turkic-speaking peoples found in all Eurasia are the result of several nomadic migrations^14,70^, which cover an area ranging from Siberia to Eastern Europe and the Middle East, through Central Asia and have been occurring during a wide period, the 5th–16th centuries^14^. In regions other than Central Asia, several studies have shown that Turkic-speaking peoples genetically resemble their geographic neighbours, with no clear genetic signal that would distinguish them^14,70^. This lends to support the model of a language replacement by elite-dominance rather than by demic diffusion for languages of the Turkish family expansion^70^. Turkmens fit in this global model but are an exception in their region. Indeed, the other Turkic-speaking populations, like Kyrgyz or Kazakhs, show a different genetic profile with a clear dominant East-Asian and Baikal components, attesting to a more significant admixture with nomads from South-Siberia and Mongolia, which have been dated around the 10^th^-14^th^ centuries^14^. The small amount of East-Asian ancestry in Turkmen has been linked to an admixture dated around the 15^th^ century, so slightly after the first admixture in Central Asia, and may come from gene-flow with these Turco-Mongol groups.

The question of the diffusion of Indo-European languages has been a hot topic in the last few years^23,50,71^. The expansion of Yamnaya related populations westward during the late Neolithic, and eastward during the Bronze Age, through the migration of Andronovo groups, suggests that they were speakers of such languages. Interestingly, the ancestry pattern found in Indo-Iranian speakers from Central Asia is not found in other Indo-Iranian speaking populations, namely, the Iranians Persians^67^. This ethnic group displays a genetic continuity since the Bronze Age with ancient individuals from Iran, with limited gene flow from the Steppes (either Central or Eastern)^67^. Furthermore, our study of the Turkmen population presents another example where language and genetics do not match, questioning the idea of inferring language displacement using population movement. Their genetic affiliation to modern Western Eurasian populations, seen in earlier studies, is due to a common steppe ancestry.

## CONCLUSION

Our results bring to light that for Indo-Iranian speakers various patterns of genetic and linguistic continuity or discontinuity coexisted through time. In southern Central Asia, we show that the actual Indo-Iranians are the product of a long-term continuity since the Iron Age with only limited recent impulses from other Eurasian groups. Our results provide further evidence that the demography of this region is complex and needs small-scale studies like this one to be fully understood. From this perspective, the precise timing of these impulses cannot be solved until more genetic data from samples from the Iron Age and historical times, who do not belong to the Steppe cultural complex, have been obtained.

## METHODS

### Compiling and merging genomic data

We selected 3102 published ancient human genomes from Eurasia (Table S1) from Paleolithic to Middle Age, whom DNA sequencing data generated with whole genome shotgun or hybridization capture technics, from the merge dataset v42.4 available at https://reich.hms.harvard.edu/allen-ancient-dna-resource-aadr-downloadable-genotypes-present-day-and-ancient-dna-data. We retained non-related individuals with more than 10,000 SNPs hit on the 1240k panel. We also added ancient individuals from two recent publications about Middle East^37^ and Mongolian Steppe^38^.

The ancient dataset was classified based on geographical, chronological, and ancestry criteria. The individuals from the Steppe with known ancestry (usually related to their localization) were labelled as *Westen_Steppe, Central_Steppe* or *Eastern_Stepppe* respectively referring to a population with a close genetic affinity with Western European Hunter Gatherer (WEHG; Loschbour, LaBrana) or Eastern European HG (EEHG; Popovo HG, Sidelkino, Karelia HG, Samara HG); to individuals with a higher ancestry from Western Siberian HG (WSHG; Tyumen Hg and Sosonivoy HG), a strong affinity with Baikal HG (BHG, Shamanka HG) or with eastern non-Africans; to populations exhibiting an East Asian component, although not all of them do like for example Andronovo population.

We analysed ancient genotypes with 1388 Eurasian individuals, 109 Yoruba, and 3 Mbuti individuals from two modern publicly available datasets: the SGDP dataset^30^, the 1000 Genomes dataset^29^. Furthermore, we also used a Central Asian specific dataset^51^ obtained in our lab using capture including 527 individuals. We merged these modern data using *mergeit* from EIGENSOFT^47^ and we haploidized them by randomly selecting one allele per position. The final merge includes 237,644 SNPs for 5129 individuals. For the analysis requiring more SNPs, we used individuals sequenced by shotgun from only three populations in the Central Asia dataset: 3 Yaghnobis (TJY), 19 Tajiks (TJE) and 24 Turkmens (TUR), and we pseudo-haploidized and combined them with the 1240k panel to obtain a second dataset of 716 743 SNPs and 4648 individuals.

### PCA

We performed PCA with *smartpca*^47^ on 1915 Eurasian present-day individuals and we projected all the 3109 ancient samples on top of the 3 best PCs. We used default parameters with *lsqproject: YES,* and *numoutlieriter: 0* settings.

### Admixture

We performed ADMIXTURE analysis^46^ on 1915 Eurasian present-day individuals and 3109 ancient samples from the first dataset on a subset of 236 665 SNPs pruned for linkage disequilibrium (by using PLINK^72^ *--indep-pairwise 200 25 0.4* function). We run ADMIXTURE a**nalyses** for clustering s**omething** K between 2 and 15, with 20 replicate **analyses** performed for each K from which we kept the most likely.

### D and F3-statistics

We computed the f3 outgroup statistics using the *qp3Pop* program with the inbreed option set to YES on our second dataset and D-statistics using the *qpDstat* program of the ADMIXTOOLS package^49^. We used Mbuti as the outgroup for both statistics in the main text, but we obtained similar results using Yoruba population as the outgroup (not shown).

### qpAdm analysis

We performed rotating *qpAdm* analysis with ADMIXTOOLS package to model the ancestry of Central Asian modern populations. For Indo-Iranians, we used Mbuti, Han, Natufian, WEHG, Ust-Ishim, MA1, Kostenki14, EEHG as *reference* populations.

Prior to the analysis, we checked if the reference populations could well discriminate between the source populations by computing f4-stat of the form f4(Mbuti, Source1, RefX, RefY), for all the sources and all the combinations of outgroups possible. Then we plotted the pairwise f4 and calculated a correlation score. We observed that our dataset discriminates well *BMAC* and *Iran_N*.

We first tested a rotating group with *Turkmenistan_IA, XiongNu,* and *GoldenHordeAsian* to assert the best source of East-Asian ancestry in our model. To test different models, we used *Iran_N, Europe_EN, BMAC, Turkmenistan_IA, XiongNu, Ukraine_Scythian* and *Germany_Corded_Ware* as rotating source populations. And to model Tajiks, we add *Indian_GreatAndaman_100BP* to represent a deep ancestry from South Asia.

To model *Turkmenistan_IA,* we used the same reference group with *Iran_N* and BHG added and we used *Germany CordedWare, Russia Poltavka, XiongNu, EuropeEN, BMAC, Andronovo, Ukraine_Scythian* as the rotating group. We also performed tests with smaller rotating groups: (1) *Andronovo, Alakul_Lisakovskiy, BMAC* (2) *Andronovo, Fedorovo_Shoindykol, BMAC* (3) *Andronovo, Sintashta, BMAC* (4) *Andronovo, Karasuk, BMAC,* (4) *Andronovo, Afanasievo, BMAC.*

To model Turkmens, we used the same reference group as for *Turkmenistan_IA,* and used TJY, *Mongolia_XiongNu, Kazakhstan_GoldenHordeAsian,* TAB, *Turkmenistan_IA* as the rotating group.

### DATES

We used *DATES* v753^18^ to estimate the time of admixture events in Tajiks, Yaghnobis, and Turkmens. To convert the estimated admixture date in generations into years, we assumed 29 years per generation^49^. The standard errors of *DATES* estimates come from the weighted block jackknife with “binsize: 0.001,” ‘‘maxdis: 1,” ‘‘runmode: 1,” ‘‘mincount: 1,” ‘‘lovalfit: 0.45” as parameters as in the example file at *https://github.com/priyamoorjani/DATES/blob/master/example/par.dates.*

## ACKNOWLEDGEMENTS

P.G.V. is supported by a PhD grant from E.N.S de Lyon. We would like to thank Ludovic Orlando, Laure Ségurel and Paul Verdu for their fruitful comments about this work. We would like to thank Romain Laurent for their assistance. We would like to thank Amanda Graham and Marcus Kearsey for proofreading the English. J.B.S addresses his acknowledgments to the Mission archéologique franco-turkmène (MAFTur), the French Ministry of Foreign Affairs (MEAE), the French Archaeological Delegation in Afghanistan (DAFA), the Shelby White and Leon Levy Program for Archaeological Publication.

## AUTHOR CONTRIBUTIONS

E.H. and C.B supervised the study. P.G.V. performed all the population genetics analyses. J.B.S. provided input about the archeology and the history of the region. N.M. processed the Central Asian dataset. P.G.V. wrote the manuscript with contributions from all co-authors.

---

**Table 1:** Plausible models for Yaghnobis (TJY), Tajiks (TJA, TAB), Turkmens (TUR) and *Turkmenistan_IA* (TurkIA) as a mixture of two or three sources obtained with *qpAdm*.

| Target | Source left populations |  |  | p-value | Admixture proportion |  |  | Standard error |  |  |
| --- | --- | --- | --- | --- | --- | --- | --- | --- | --- | --- |
|  | A | B | C |  | A | B | C | A | B | C |
| TJY | TurkmenIA | XiongNu |  | 0.2462 | 0.933 | 0.067 |  | 0.010 | 0.010 |  |
| TJY | TurkmenIA | XiongNu | EuropeEN | 0.2691 | 0.897 | 0.071 | 0.032 | 0,044 | 0.010 | 0.041 |
| TJY | TurkmenIA | XiongNu | BMAC | 0.1301 | 0.906 | 0.068 | 0.026 | 0.104 | 0.011 | 0.107 |
| TJY | TurkmenIA | XiongNu | UkraineScythian | 0.0911 | 0.899 | 0.069 | 0.032 | 0.052 | 0.010 | 0.048 |
| TJA | TurkmenIA | XiongNu | GreatAndaman_100BP | 0.2780 | 0.709 | 0.165 | 0.129 | 0.053 | 0.022 | 0.072 |
| TUR | TAB | GoldenHorde |  | 0.48007 | 0.941 | 0.059 |  | 0.005 | 0.005 |  |
| TurkIA | BMAC | Andronovo |  | 0.327016 | 0.471 | 0.529 |  | 0.036 | 0.036 |  |

## Notes

### Competing Interest Statement

The authors have declared no competing interest.

## References

1. Wells, R. S. et al. The Eurasian Heartland: A continental perspective on Y-chromosome diversity. Proc. Natl. Acad. Sci. 98, 10244–10249 (2001).

2. Martínez-Cruz, B. et al. In the heartland of Eurasia: the multilocus genetic landscape of Central Asian populations. Eur. J. Hum. Genet. 19, 216–223 (2011).

3. Brunet, F., Tengberg, M. & Harris, D. R. Origins of Agriculture in Western Central Asia. An Environmental-Archaeological Study. Philadelphia: University of Pennsylvania Museum of Archaeology and Anthropology. Paléorient 39, 209/211 (2013).

4. Luneau, E. L’âge du bronze final en Asie centrale méridionale (1750-1500/1450 avant notre ère: la fin de la civilisation de l’Oxus). (UNIVERSITÉ PARIS 1 - PANTHÉON-SORBONNE, 2010).

5. Bendezu-Sarmiento, J. & Lhuillier, J. Transitions socioculturelles lors de la fin de la civilisation de l’Oxus. Les Nouv. l’archéologie 32–40 (2020). doi:10.4000/nda.10466

6. Bonora, G. L. The Oxus Cilization and the Northern Steppes. in The World of the Oxus Civilization (eds. Lyonnet, B. & Dubova, N.) 734–775 (Routledge, 2021).

7. Bendezu-Sarmiento, J. & Lhuillier, J. Sine Sepulchro cultural complex of Transoxiana (between 1500 and the middle of the 1st Millennium BCE). Funerary Practices of the Iron Age in Southern Central Asia: Recent Work, old Data, and new Hypotheses. Archäologische Mitteilungen aus Iran und Turan 45, 281–315 (2015).

8. Luneau, E. Les mutations sociopolitiques de la période finale de la civilisation de l’Oxus. in Les marqueurs archéologiques du pouvoir (eds. Brunet, O., Sauvin, C.-E. & Halabi, T.) 221–241 (2013).

9. Lhuillier, J. The settlement pattern in Central Asia during the Early Iron Age. in Urban cultures of Central Asia from the Bronze Age to the Karakhanids. Learnings and conclusions from new archaeological investigations and discoveries, Proceedings of the First International Congress on Central Asian Archaeology held at the University of (ed. C. Baumer, M. N.) 12, 115–128 (Harrassowitz Verlag, 2019).

10. Krader, L. Peoples of Central Asia, 2nd edition. (Indiana University Publications, Bloomington, 1966).

11. Palstra, F. P., Heyer, E. & Austerlitz, F. Statistical Inference on Genetic Data Reveals the Complex Demographic History of Human Populations in Central Asia. Mol. Biol. Evol. 32, 1411–1424 (2015).

12. Cilli, E. et al. The genetic legacy of the Yaghnobis: A witness of an ancient Eurasian ancestry in the historically reshuffled central Asian gene pool. Am. J. Phys. Anthropol. 168, 717–728 (2019).

13. Damgaard, P. de B. et al. 137 ancient human genomes from across the Eurasian steppes. Nature 557, 369–374 (2018).

14. Yunusbayev, B. et al. The genetic legacy of the expansion of Turkic-speaking nomads across Eurasia. PLoS Genet. 11, e1005068 (2015).

15. Marchi, N. et al. Sex-specific genetic diversity is shaped by cultural factors in Inner Asian human populations. Am. J. Phys. Anthropol. 162, 627–640 (2017).

16. Heyer, E. & Mennecier, P. Genetic and linguistic diversity in Central Asia. in Becoming eloquent (eds. D’Errico, F. & Hombert, J.-M.) 163–180 (John Benjamins Publishing Company, 2009).

17. de Barros Damgaard, P. et al. The first horse herders and the impact of early Bronze Age steppe expansions into Asia. Science (80-.). 360, eaar7711 (2018).

18. Narasimhan, V. M. et al. The formation of human populations in South and Central Asia. Science (80-.). 365, eaat7487 (2019).

19. Järve, M. et al. Shifts in the Genetic Landscape of the Western Eurasian Steppe Associated with the Beginning and End of the Scythian Dominance. Curr. Biol. 29, 2430–2441 (2019).

20. Saag, L. et al. The Arrival of Siberian Ancestry Connecting the Eastern Baltic to Uralic Speakers further East. Curr. Biol. 29, 1701–1711.e16 (2019).

21. Olalde, I. et al. The genomic history of the Iberian Peninsula over the past 8000 years. Science (80-.). 363, 1230–1234 (2019).

22. Olalde, I. et al. The Beaker phenomenon and the genomic transformation of northwest Europe. Nature 555, 190–196 (2018).

23. Haak, W. et al. Massive migration from the steppe was a source for Indo-European languages in Europe. Nature 522, 207–211 (2015).

24. Mathieson, I. et al. Genome-wide patterns of selection in 230 ancient Eurasians. Nature 528, 499–503 (2015).

25. Stolarek, I. et al. Goth migration induced changes in the matrilineal genetic structure of the central-east European population. Sci. Rep. 9, 6737 (2019).

26. Mathieson, I. et al. The genomic history of southeastern Europe. Nature 555, 197–203 (2018).

27. Unterländer, M. et al. Ancestry and demography and descendants of Iron Age nomads of the Eurasian Steppe. Nat. Commun. 8, 14615 (2017).

28. Krzewińska, M. et al. Ancient genomes suggest the eastern Pontic-Caspian steppe as the source of western Iron Age nomads. Sci. Adv. 4, eaat4457 (2018).

29. The 1000 Genomes Project Consortium et al. A global reference for human genetic variation. Nature 526, 68–74 (2015).

30. Mallick, S. et al. The Simons Genome Diversity Project: 300 genomes from 142 diverse populations. Nature 538, 201–206 (2016).

31. Fu, Q. et al. The genetic history of Ice Age Europe. Nature 534, 200–205 (2016).

32. Wang, C.-C. et al. Ancient human genome-wide data from a 3000-year interval in the Caucasus corresponds with eco-geographic regions. Nat. Commun. 10, 590 (2019).

33. Broushaki, F. et al. Early Neolithic genomes from the eastern Fertile Crescent. Science (80-.). 353, 499–503 (2016).

34. Lazaridis, I. P. et al. Genomic insights into the origin of farming in the ancient Near East. Nature 536, 419–424 (2016).

35. Fu, Q. et al. Genome sequence of a 45,000-year-old modern human from western Siberia. Nature 514, 445–449 (2014).

36. Vinet, L. & Zhedanov, A. A ‘missing’ family of classical orthogonal polynomials. Proc. Natl. Acad. Sci. 113, 6886–6891 (2010).

37. Skourtanioti, E. et al. Genomic History of Neolithic to Bronze Age Anatolia, Northern Levant, and Southern Caucasus. Cell 181, 1158–1175.e28 (2020).

38. Jeong, C. et al. A Dynamic 6,000-Year Genetic History of Eurasia’s Eastern Steppe. Cell 183, 890–904.e29 (2020).

39. Sikora, M. et al. Ancient genomes show social and reproductive behavior of early Upper Paleolithic foragers. Science (80-.). 358, 659–662 (2017).

40. Sikora, M. et al. The population history of northeastern Siberia since the Pleistocene. Nature 570, 182–188 (2019).

41. Raghavan, M. et al. Upper Palaeolithic Siberian genome reveals dual ancestry of Native Americans. Nature 505, 87–91 (2014).

42. Mittnik, A. et al. The genetic prehistory of the Baltic Sea region. Nat. Commun. 9, 442 (2018).

43. Veeramah, K. R. et al. Population genomic analysis of elongated skulls reveals extensive female-biased immigration in Early Medieval Bavaria. Proc. Natl. Acad. Sci. 115, 3494–3499 (2018).

44. Jeong, C. et al. Bronze Age population dynamics and the rise of dairy pastoralism on the eastern Eurasian steppe. Proc. Natl. Acad. Sci. 115, E11248–E11255 (2018).

45. Jeong, C. et al. Long-term genetic stability and a high-altitude East Asian origin for the peoples of the high valleys of the Himalayan arc. Proc. Natl. Acad. Sci. 113, 7485–7490 (2016).

46. Alexander, D. H., Novembre, J. & Lange, K. Fast model-based estimation of ancestry in unrelated individuals. Genome Res. 19, 1655–64 (2009).

47. Patterson, N., Price, A. L. & Reich, D. Population Structure and Eigenanalysis. PLoS Genet. 2, e190 (2006).

48. Moreno-Mayar, J. V. et al. Terminal Pleistocene Alaskan genome reveals first founding population of Native Americans. Nature 553, 203–207 (2018).

49. Patterson, N. et al. Ancient Admixture in Human History. Genetics 192, 1065–1093 (2012).

50. Allentoft, M. E. et al. Population genomics of Bronze Age Eurasia. Nature 522, 167–172 (2015).

51. Marchi, N. et al. Close inbreeding and low genetic diversity in Inner Asian human populations despite geographical exogamy. Sci. Rep. 8, (2018).

52. Heyer, E. et al. Genetic diversity and the emergence of ethnic groups in Central Asia. BMC Genet. 10, 49 (2009).

53. Chaix, R., Austerlitz, F., Hegay, T., Quintana-Murci, L. & Heyer, E. Genetic traces of east-to-west human expansion waves in Eurasia. Am. J. Phys. Anthropol. 136, 309–317 (2008).

54. Grousset, R. L’empire des steppes. Attila, Gengis-khan, Tamerla. (1965).

55. Brunet, F. La néolithisation en Asie centrale: un état de la question. Paléorient 24, 27–48 (1998).

56. Bendezu-Sarmiento, J., Ismagulova, A., Bajpakov, K. & Samashev, Z. De l’âge du Bronze à l’âge du Fer au Kazakhstan, gestes funéraires et paramètres biologiques. Identités culturelles des populations Andronovo et Saka. in Mémoires de la Mission Archéologique Française en Asie centrale, T. XII 606 (2007).

57. Rasmussen, S. et al. Early Divergent Strains of Yersinia pestis in Eurasia 5,000 Years Ago. Cell 163, 571–582 (2015).

58. Gnecchi-Ruscone, G. A. et al. Ancient genomic time transect from the Central Asian Steppe unravels the history of the Scythians. Sci. Adv. 7, eabe4414 (2021).

59. Barbara Cerasetti. WHO INTERACTED WITH WHOM? Redefining the interaction between BMAC people and mobile pastoralists in Bronze Age southern Turkmenistan. in The World of the Oxus Civilization (eds. Lyonnet, B. & Dubova, N. A.) 487–495 (Routledge, 2021).

60. Luneau, E. The End of the Oxus Civilization. in The World of the Oxus Civilization (eds. Lyonnet, B. & Dubova, N.) 496–524 (Routledge, 2021).

61. Masson, V. M. History of Civilisations of Central Asia Volume 1. (UNESCO, 1992).

62. A Millennium of History The Iron Age in southern Central Asia (2nd and 1st Millennia BC). Proceedings of the conference held in Berlin (June 23-25, 2014). Dedicated to the memory of Viktor Ivanovich Sarianidi. (DIETRICH REIMER VERLAG, BERLIN, 2018).

63. Zhabagin, M. et al. The Connection of the Genetic, Cultural and Geographic Landscapes of Transoxiana. Sci. Rep. 7, 3085 (2017).

64. Novák, L’. Jaghnóbsko-český slovník s přehledem jaghnóbské gramatiky. (Filozofická fakulta Karlovy univerzity v Praze, 2010).

65. Thouzeau, V., Mennecier, P., Verdu, P. & Austerlitz, F. Genetic and linguistic histories in Central Asia inferred using approximate Bayesian computations. Proc. R. Soc. B Biol. Sci. 284, 20170706 (2017).

66. Li, H., Cho, K., Kidd, J. R. & Kidd, K. K. Genetic Landscape of Eurasia and “Admixture” in Uyghurs. Am. J. Hum. Genet. 85, 934–937 (2009).

67. Mehrjoo, Z. et al. Distinct genetic variation and heterogeneity of the Iranian population. PLOS Genet. 15, e1008385 (2019).

68. Jarrige, J.-F. Les relations archéologiques entre les régions au sud et au nord de l’Hindu Kush du Ve millénaire jusqu’au milieu du IIIe millénaire avant notre ère à la lumière des données fournies par les sites de la région de Kachi-Bolan au Balochistan pakistanais. Cah. d’Asie Cent. 41–68 (2013).

69. Lhuillier, J. What about the relationships between the sites with painted pottery North and South of Hindu-Kush during the transition from Bronze Age to Early Iron Age? Reassessment of data and new perspectives. in South Asian Archaeology and Art 2012 1, 155–168 (Brepols, 2012).

70. Triska, P. et al. Between Lake Baikal and the Baltic Sea: genomic history of the gateway to Europe. BMC Genet. 18, 110 (2017).

71. Klejn, L. S. et al. Discussion: Are the origins of indo-european languages explained by the migration of the Yamnaya Culture to the West? Eur. J. Archaeol. 21, 3–17 (2018).

72. Vinet, L. & Zhedanov, A. A ‘missing’ family of classical orthogonal polynomials. Am. J. Hum. Genet. 81, 559–575 (2010).

