## Supplementary material for "Genetic Continuity of Indo-Iranian Speakers Since the Iron Age in Southern Central Asia"

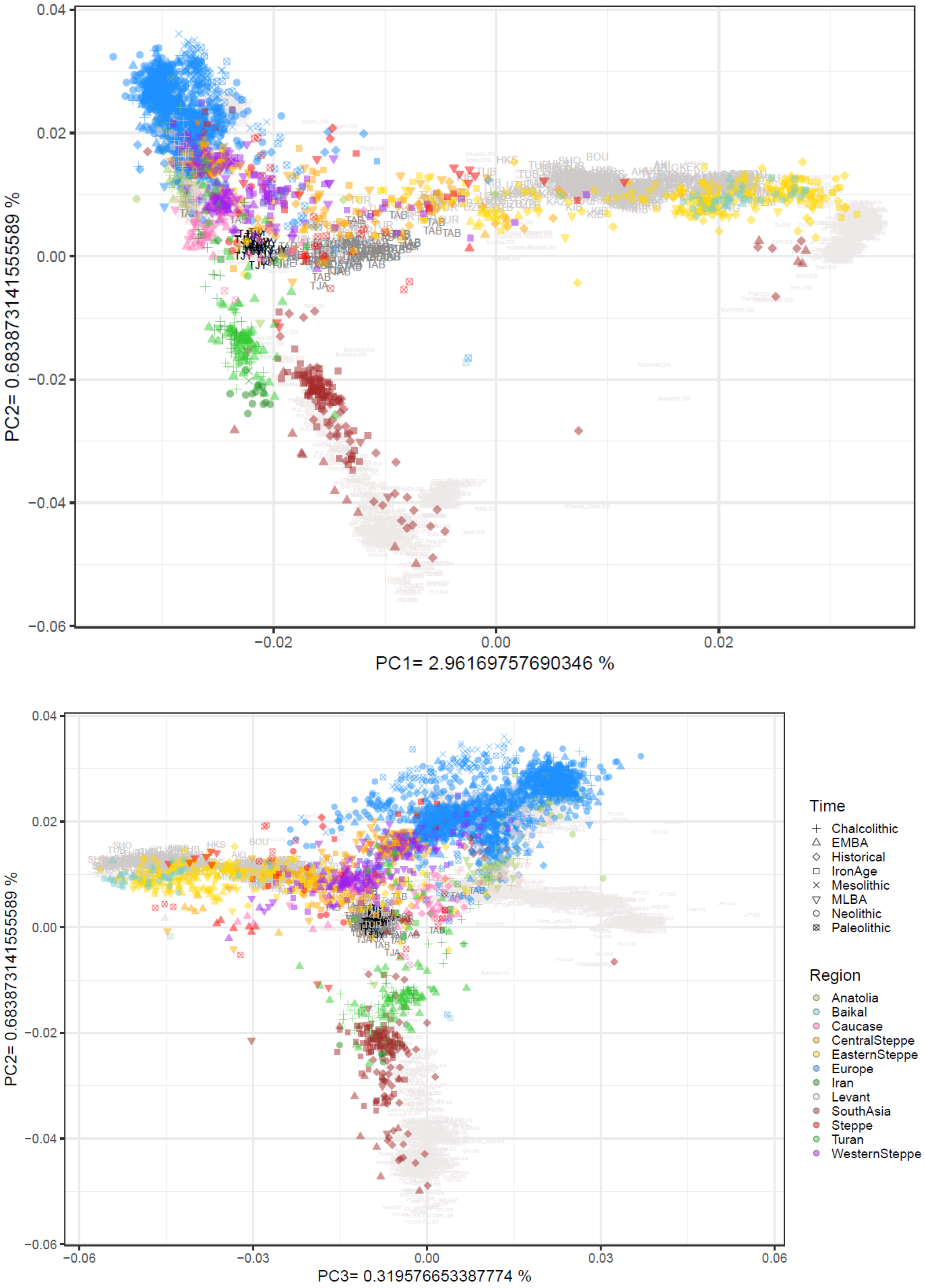


**Figure S1. Full PCA** computed on 1915 modern genomes, with projection of 3102 ancient genomes

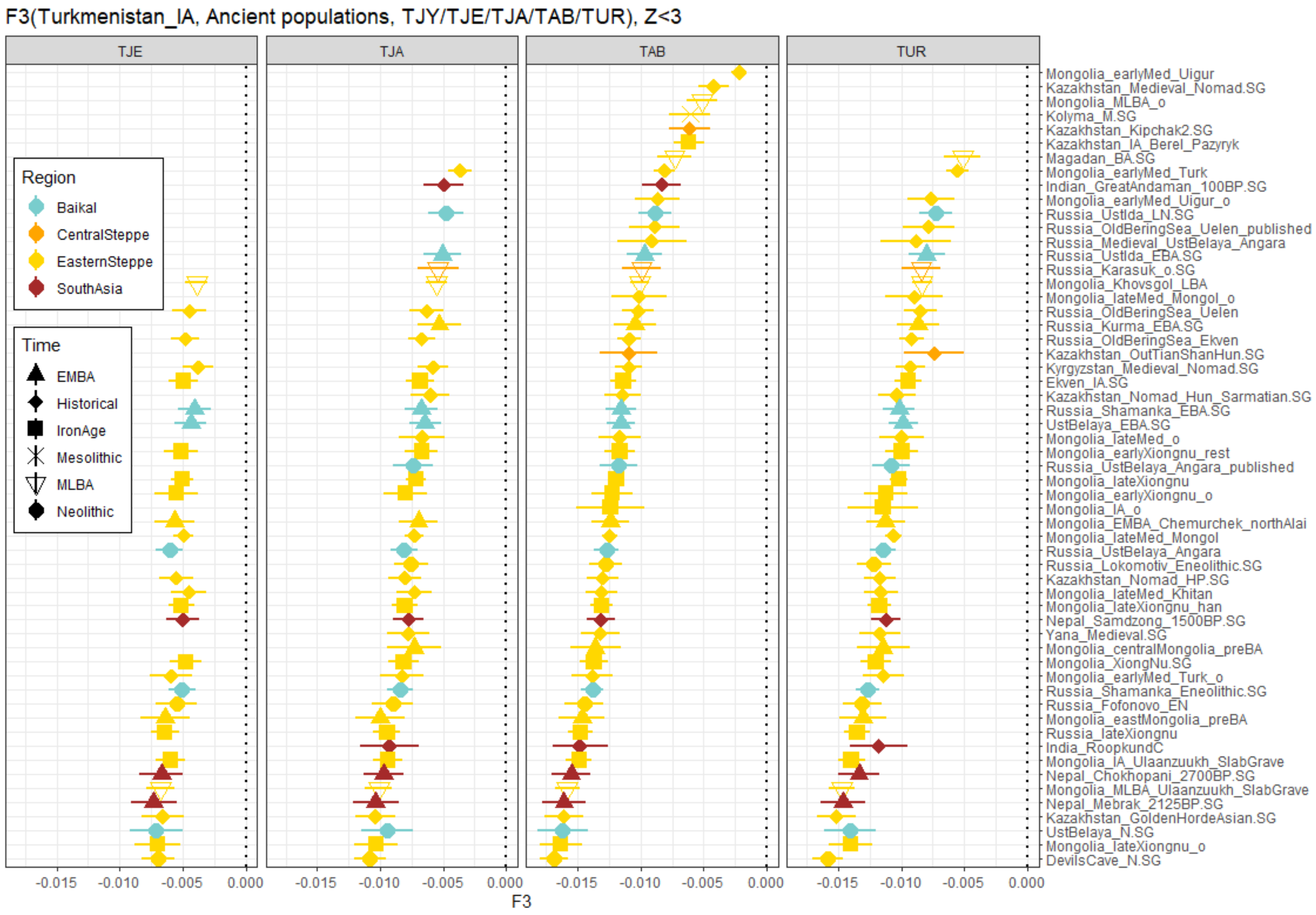


**Figure S2. All the significatively negative admixture f3-statistics** in the form f3(*Turkmenistan_IA*, *Ancient population*, TJY/TJE/TJA/TAB/TUR) calculated on the first dataset. We did not obtain significative f3 for TJY.

**Figure S3 – Full ADMIXTURE analysis for all ancient individuals**

**Figure S4 – Full ADMIXTURE analysis for all present-day individuals**

**Table S1 – Contextual information for published ancient genome-wide data**

**Table S2 – Contextual information for published modern genomes from all Eurasia**

**Table S3 – Contextual information for modern genome-wide data from Central Asia (300k SNPs array)**

**Table S4 – Contextual information for modern genome from Central Asia (Shotgun)**

**Table S5 – All D-stat of the form D(Mbuti, Ancient pop; Iron Age Turan, TAB/TJA/TJE/TJY/TUR)**

**Table S6 – All f3-admixture of the form f3(Ancient pop, Iron Age Turan; TAB/TJA/TJE/TJY/TUR)**

**Table S7 - Significantly negative f3-admixture of the form f3(Ancient Pop1, Ancient Pop2; TJY/TJE/TJA/TAB)**

**Table S8 – All f3-outgroup of the form f3(Ancient pop, Central Asia pop; Outgroup)**
